# Translational arrest and mRNA decay are independent activities of alphaherpesvirus virion host shutoff proteins

**DOI:** 10.1101/2024.02.02.578636

**Authors:** Lucy Eke, Alistair Tweedie, Sophie Cutts, Emma L Wise, Gillian Elliott

## Abstract

The herpes simplex virus 1 (HSV1) virion host shutoff (vhs) protein is an endoribonuclease that regulates the translational environment of the infected cell, by inducing the degradation of host mRNA via cellular exonuclease activity. To further understand the relationship between translational shutoff and mRNA decay, we have used ectopic expression to compare HSV1 vhs (vhsH) to its homologues from four other alphaherpesviruses – varicella zoster virus (vhsV), bovine herpesvirus 1 (vhsB), equine herpesvirus 1 (vhsE) and Marek’s disease virus (vhsM). Only vhsH, vhsB and vhsE induced degradation of a reporter luciferase mRNA, with polyA+ *in situ* hybridisation indicating a global depletion of cytoplasmic polyA+ RNA and a concomitant increase in nuclear polyA+ RNA and the polyA tail binding protein PABPC1 in cells expressing these variants. By contrast, vhsV and vhsM failed to induce reporter mRNA decay and polyA+ depletion, but rather, induced cytoplasmic G3BP1 and polyA+ mRNA-containing granules and phosphorylation of the stress response proteins eIF2α and protein kinase R. Intriguingly, regardless of their apparent endoribonuclease activity, all vhs homologues induced an equivalent general blockade to translation as measured by single cell puromycin incorporation. Taken together, these data suggest that the activities of translational arrest and mRNA decay induced by vhs are separable and we propose that they represent sequential steps of the vhs host interaction pathway.

## Introduction

A wide range of viruses, including influenza viruses, coronaviruses and herpesviruses express endoribonucleases to help regulate the mRNA environment in the infected cell and favour virus over host translation (1–6). In the case of the alphaherpesvirus herpes simplex virus 1 (HSV1), the product of the UL41 gene, named the virion host shutoff protein (vhs), is well documented as an endoribonuclease that functions not only during infection (7–9), but also when expressed in isolation, where it has been shown to abrogate the expression of reporter proteins such as beta-galactosidase and luciferases (10–14). vhs is proposed to induce the degradation of mRNA (15–17) by binding to the cellular translation initiation machinery through the eIF4A and eIF4H components of the eIF4F cap-binding complex, resulting in mRNA cleavage (18–21). It is hypothesized that as for the KSHV SOX protein, the cellular exonuclease XRN1 is subsequently recruited to cleave mRNA and accelerate its decay in the cytoplasm (22). An additional consequence of vhs-induced mRNA decay is the accumulation of polyA binding protein (PABPC1) in the nucleus (13, 23, 24). Ordinarily, PABPC1 cycles between the nucleus and cytoplasm on the polyA tails of exporting mRNAs exhibiting a steady state cytoplasmic localisation (25, 26). When this balance is disturbed through enhanced mRNA decay in the cytoplasm, PABPC1 accumulates in the nucleus along with other RNA binding proteins (27) resulting in the hyperadenylation of mRNA and a subsequent block to mRNA export (23, 26). Recently, vhs has also been shown to induce the degradation of double-stranded RNA contributing to the inhibition of the cellular protein kinase R (PKR) response seen in virus infection (28), and providing an explanation for the documented induction of stress granules in cells infected with vhs knockout viruses (29, 30). It is suggested that in the absence of vhs, dsRNA forms as a consequence of transcription of both strands of the HSV1 genome, inducing the phosphorylation of PKR and eIF2α, the stalling of translation and induction of stress granules (31), thereby shutting down viral as well as cellular protein translation.

The UL41 gene is present in all alphaherpesviruses, and alignment of the proteins has previously identified four conserved domains across the protein (32). In HSV1 vhs, these regions (Fig 1) have been shown to be important for vhs activity (12, 33). The activity of the other alphaherpesvirus vhs proteins is poorly defined in comparison to HSV1 vhs: homologues from pseudorabies virus and bovine herpesvirus 1 (BHV1) have been shown to function when expressed either ectopically or in the virus (34–38); equine herpesvirus 1 (EHV1) and Marek’s disease virus (MDV) homologues appear to be active when expressed in isolation but not in the context of virus infection (39–41); and there are conflicting reports on the functionality of varicella zoster virus (VZV) vhs (42, 43). As the level of identity between these proteins ranges from only 33 to 51% (Table 1), there is scope for differential activity amongst the homologues.

**Figure 1:**
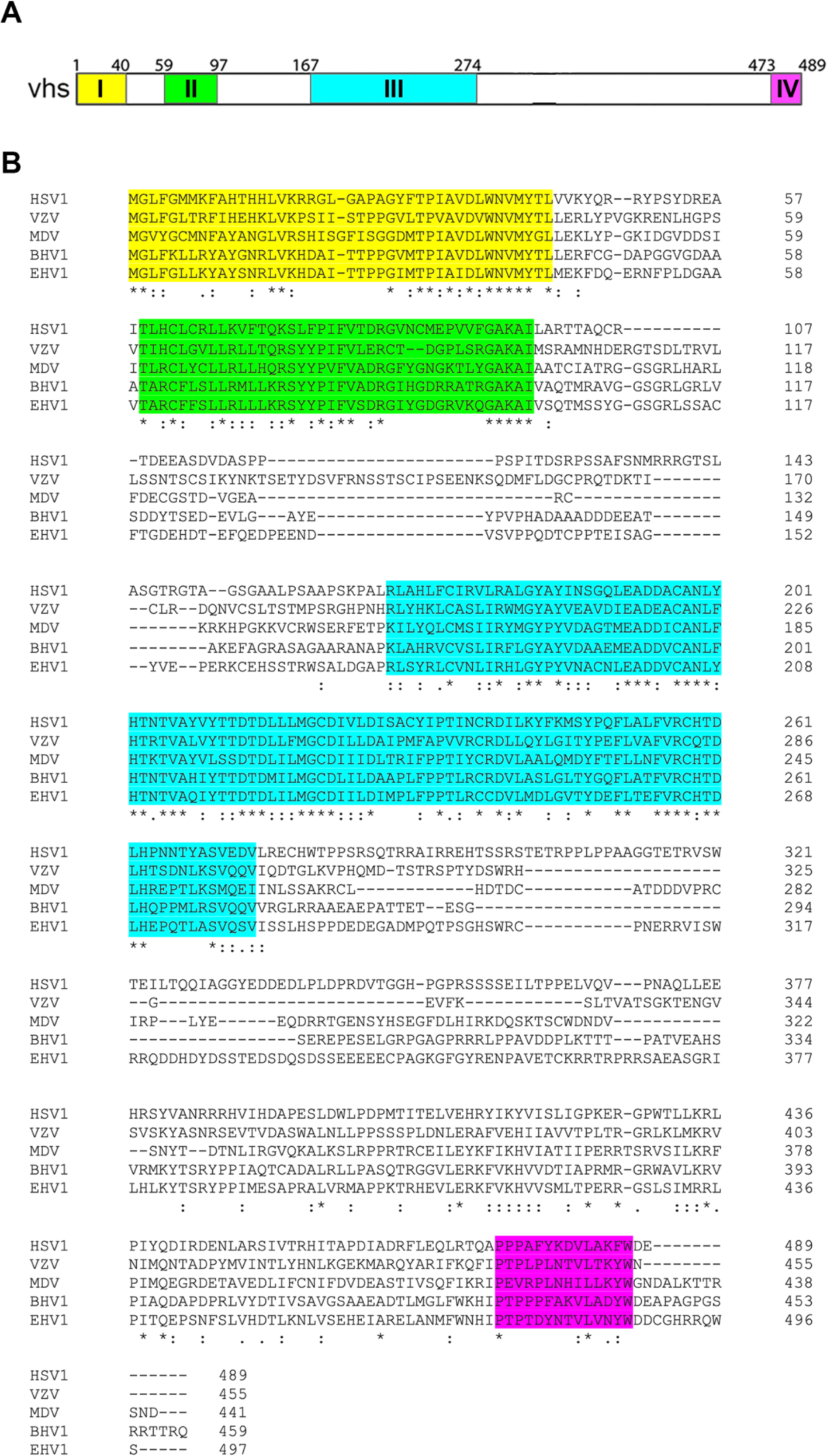
Amino acid alignment of vhs homologues from HSV1, VZV, BHV1, EHV1 and MDV. **(A)** Line drawing of the HSV1 vhs open reading frame. Four conserved areas are coloured and based on (32). **(B)** A multiple sequence alignment for vhs homologues from HSV1 (YP_009137116), VZV (NP_040140), MDV (YP_001033970), BHV1 (NP_045317) and EHV1 (YP_053064) was performed using Clustal Omega with default settings.

**Table 1.** Identity matrix between the homologues of vhs used in this study.

|  | HSV1 | VZV | MDV | BHV1 | EHV1 |
| --- | --- | --- | --- | --- | --- |
| HSV1 | 100 | 33.57 | 33.01 | 35.75 | 33.48 |
| VZV | 33.57 | 100 | 38.44 | 40.77 | 40.46 |
| MDV | 33.01 | 38.44 | 100 | 41.35 | 39.82 |
| BHV1 | 35.75 | 40.77 | 41.35 | 100 | 51.1 |
| EHV1 | 33.48 | 40.46 | 39.82 | 51.1 | 100 |

Here we have evaluated the relative activity of VZV (vhsV), BHV1 (vhsB), EHV1 (vhsE) and MDV (vhsM) homologues in relation to that from HSV1 (vhsH) and have compared their behaviour when expressed in isolation of virus infection. We show that vhsB and vhsE behave in a similar fashion to vhsH, abrogating gene expression from a reporter plasmid and inducing the degradation of its mRNA. They also induce a general depletion of cytoplasmic polyadenylated (polyA+) RNA and the relocalisation of PABPC1 to the nucleus; of note, vhsB was seen to be the most highly active vhs homologue. By contrast, vhsV does not degrade the reporter mRNA or deplete the cytoplasm of polyA+ RNA, but instead causes the formation of G3BP1-containing RNA granules in expressing cells; vhsM does not degrade mRNA but exhibits PABPC1 relocalisation characteristics of both vhsH and vhsV. Moreover, despite the lack of mRNA decay in cells expressing vhsV or vhsM, single cell puromycin incorporation revealed that all five vhs homologues were able to induce a general blockade to cellular translation, suggesting that translational arrest is independent of mRNA decay. These data highlight the benefit of studying the activity of a virus protein from different family members to unravel important mechanistic detail.

## Materials and Methods

### Cells and plasmids

Hela cells were cultured in DMEM supplemented with 10% foetal bovine serum (Invitrogen). Plasmids pEGFPC1 and pEGFPN1, expressing GFP under the control of the hCMV immediate-early promoter were obtained from Clontech. The construction of plasmids expressing vhs from HSV1 strain17 as vhs-GFP and untagged pcvhs have been described previously (13, 44). vhsB-GFP, vhsE-GFP and vhsM-GFP were constructed by PCR amplification of the following; UL41 from BHV1 strain P8-2; ORF 19 from EHV1 strain Ab4; and 054 from MDV strain RB1B followed by insertion into pEGFPN1. The VZV ORF17 gene was synthesized from reference sequence MH709372.1 and inserted into pEGFPN1. Untagged versions were constructed by replacing the GFP open reading frame with a stop codon.

### Antibodies

Commercial antibodies used in this study were α-tubulin (Sigma), GFP (Clontech), PABPC1 (Santa Cruz) G3BP1 (Sigma) and Puromycin (EMD Millipore).

### Quantitative RT-PCR (RT-qPCR)

Total RNA was extracted from cells using Qiagen RNeasy kit. Excess DNA was removed by incubation with DNase I (Invitrogen) for 15 min at room temperature, followed by inactivation for 10 min at 65°C in 25 nM of EDTA. Superscript III (Invitrogen) was used to synthesise cDNA using random primers according to manufacturer’s instructions. All RT-qPCR assays were carried out in 96-well plates using MESA Blue qPCR MasterMix Plus for SYBR Assay (Eurogentec). GLuc was amplified using GLuc forward primer CCTACGAAGGCGACAAAGAG and reverse primer TTGTGCAGTCCACACACAGA and results were normalised by amplifying 18s primer sequences. Cycling was carried out in a Lightcycler (Roche), and GLuc mRNA levels determined by the ΔΔCt method using 18s RNA as reference gene (18s F: CCAGTAAGTGCGGGTCATAAGC; 18s R: GCCTCACTAAACCATCCAATCGG).

### SDS-PAGE and Western blotting

Protein samples were analysed by SDS-polyacrylamide gel electrophoresis and transferred to nitrocellulose membrane for Western blot analysis. IRDye secondary antibodies were obtained from LI-COR and Western blots were imaged on a LI-COR Odyssey Imaging system.

### Immunofluorescence

Cells for immunofluorescence were grown on coverslips and fixed with 4% paraformaldehyde in PBS for 20 min at room temperature, followed by permeabilisation with 0.5% Triton-X100 for 10 min. Fixed cells were blocked by incubation in PBS with 10% newborn calf serum for 20 min, before the addition of primary antibody in PBS with 10% serum, and a further 30-min incubation. After extensive washing with PBS, the appropriate Alexafluor conjugated secondary antibody was added in PBS with 10% serum and incubated for a further 15 min. The coverslips were washed extensively in PBS and mounted in Mowiol containing DAPI to stain nuclei. Images were acquired using a Nikon A1 confocal microscope and processed using ImageJ software.

### Fluorescent *in situ* hybridisation (FISH) of RNA

HeLa cells were grown in 2-well slide chambers (Fisher Scientific) and transfected with plasmid. Sixteen hours later, cells were fixed for 20 min in 4% PFA. For polyA+ RNA FISH, cold methanol was added to cells for 10 mins followed by 70% ethanol for 10 mins, then 1M Tris (pH 8) for 5 mins. The cells were then incubated with 1 ng/μl 5’-labelled Cy3-oligodT in hybridization buffer (0.005% bovine serum albumin; 1 mg/ml yeast tRNA; 10% dextran sulfate; 25% deionised formamide; in 2 x SSC) at 37 °C for 2 hours before sequential washing in 4 x SSC, and 2 x SSC; 0.1% tritonX100 in 2 x SSC. Cells were mounted in Mowiol containing DAPI to stain nuclei, images acquired with a Nikon A2 inverted confocal microscope and processed using Adobe Photoshop software.

### *Gaussia* luciferase reporter assay

HeLa cells were transfected with plasmid pCMV-GLuc-1 and increasing amounts of vhs-expressing plasmids. After 16 h, the medium was replaced, sampled 4 h later, and chemiluminescence measured by injection of coelenterazine at 1 μg/ml in PBS and read on a Clariostar plate reader.

### Puromycin incorporation assay

HeLa cells were grown on coverslips before transfection with vhs-expressing plasmids. After 16 h, the medium was removed and replaced with medium containing 10 ug/ml puromycin (Invivogen) and incubated for 5 minutes at 37°C. Following incubation, cells were washed once in PBS before fixation for 20 minutes in 4 % PFA and processing for immunofluorescence.

## Results

### Confirmation of expression of five alphaherpesvirus vhs homologues

To compare the relative expression of five vhs proteins, expression vectors for the vhs open reading frames from HSV1 (vhsH), VZV (vhsV), BHV1 (vhsB), EHV1 (vhsE) and MDV (vhsM), fused at their C-termini to GFP, were constructed as a means of confirming their expression by transient transfection. Of note, we and others have shown that vhsH is expressed at very low levels in transient systems (13, 45) while fusion of vhsH to GFP confers this reduced level of expression to GFP (44). We hypothesize that this is due to a combination of a previously identified inhibitory sequence in the vhsH transcript, which blocks translation (13), and the negative feedback of vhsH endoribonuclease activity on its own transcript (13). Expression and localisation of the vhs homologues were compared by transfecting plasmids expressing the GFP-fusion proteins into HeLa cells and analysing 16 h later by SDS-PAGE and Western blotting for GFP, or by imaging of GFP fluorescence. As shown previously (13, 44), fusion of vhsH to GFP drastically reduced the expression of GFP as demonstrated by Western blotting (Fig 2A). Likewise, fusion of the other homologues to GFP reduced the expression of GFP to varying degrees, with vhsV-GFP barely detectable by Western blotting, but vhsB-GFP, vhsE-GFP and vhsM-GFP detectable, but still greatly reduced compared to GFP alone (Fig 2A). Nonetheless, despite this low level of expression, all homologue GFP fusion proteins were detected at the single cell level and were localised predominantly to the cytoplasm (Fig 2B).

**Figure 2.**
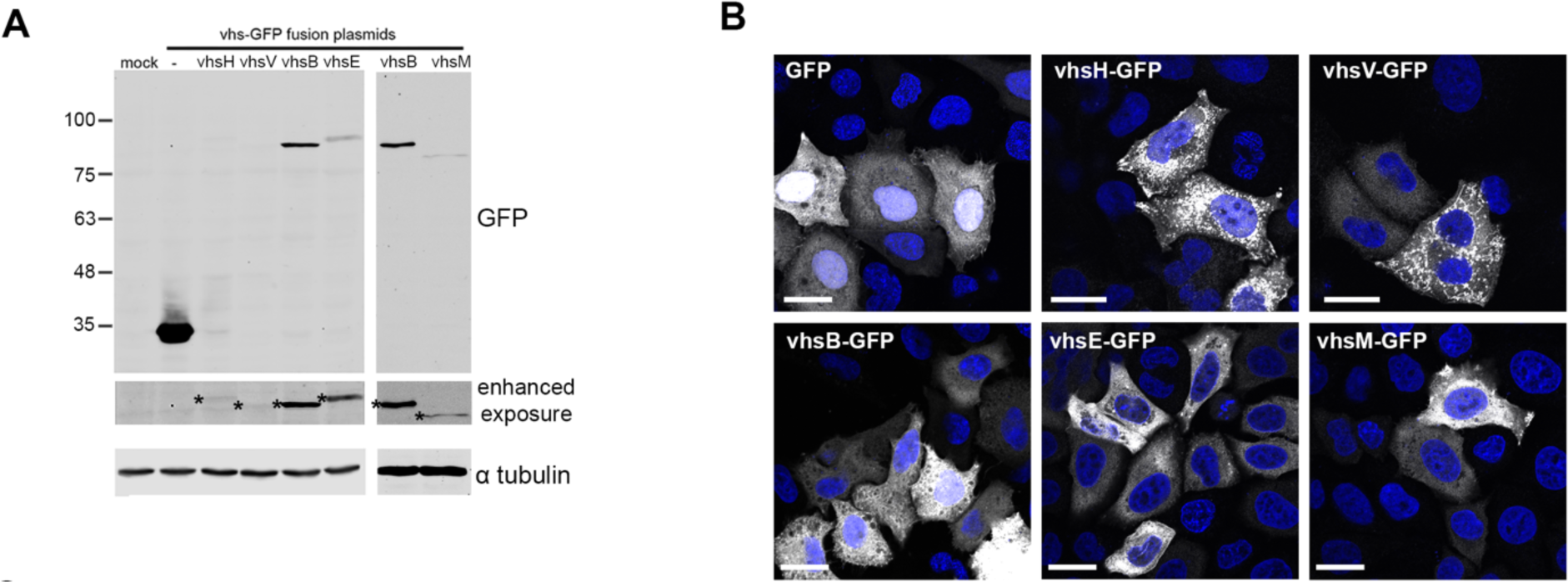
Expression of five vhs homologues as GFP fusion proteins. **(A)** HeLa cells were transfected with plasmids expressing GFP (-) or each of the vhs homologues fused at their C-termini to GFP. After 16 h total cell extracts were prepared and analysed by SDS-PAGE and Western blotting for GFP or α-tubulin as a loading control. Protein molecular weight markers are shown on left hand side (kDa). **(B)** HeLa cells grown on coverslips were transfected as for (A), fixed after 16 h and nuclei stained with DAPI (blue). GFP fluorescence (white) was imaged using a NikonA1 confocal microscope. Scale bar = 20 μm.

### Differential regulation of a GLuc reporter by the vhs homologues

Ectopic expression of vhsH is known to reduce expression of a number of reporter proteins including *Gaussia* luciferase (GLuc), a secreted reporter luminescent protein (13). As our previous work has revealed that tagging vhsH with even a small epitope can abrogate its activity, the GFP open reading frame was next removed from the vhs-GFP expression vectors above and replaced with a stop codon. These plasmids, now expressing untagged vhs, were transfected into HeLa cells together with the GLuc expressing plasmid. As a negative control, a vhsH variant (vhs*) that has a point mutation in the open reading frame known to severely attenuate its activity (13) was also included. The resulting levels of secreted GLuc indicated that the vhs homologues divided into two groups – one that comprises vhsH, vhsB and vhsE which were highly active for GLuc downregulation compared to vhs*, and one that comprises vhsV and vhsM which had only modest GLuc downregulation compared to vhs* (Fig 3A).

**Figure 3:**
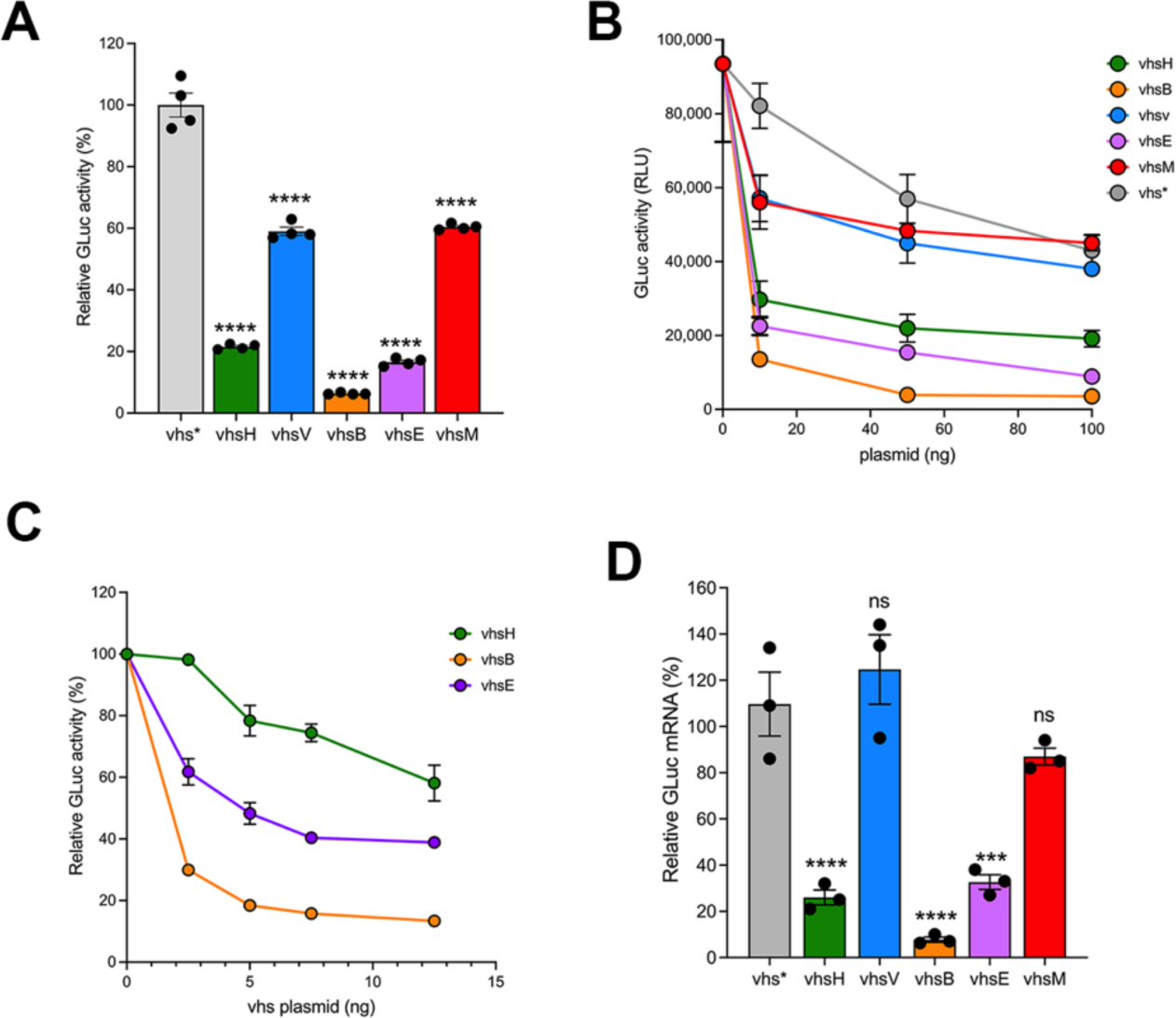
Differential effect of vhs homologues on reporter gene expression. **(A)** HeLa cells were transfected with GLuc alone or in combination with untagged vhs homologue encoding plasmids. After 16 h, media was changed and 4 h later samples were removed and analysed for GLuc activity in a Clariostar plate reader. Statistical analysis was carried out using ordinary one-way ANOVA multiple comparisons and comparison made to vhs* values. **** p < 0.0001. **(B)** HeLa cells were transfected with GLuc alone or with increasing amounts of plasmids encoding vhs*, vhsH, vhsV, vhsB, vhsE or vhsM. GLuc activity was measured as in (A). **(C)** HeLa cells were transfected with GLuc alone or with increasing amounts of plasmids encoding vhsH, vhsB or vhsE. GLuc activity was measured as in (A). **(D)** HeLa cells were transfected as for (A), and after 20 h cells total RNA was harvested and RTqPCR carried out with primers specific for GLuc, using 18s as reference gene. Differential GLuc expression was determined to GLuc expression in vhs* expressing cells using the 1ΔΔCt method. Statistical analysis was carried out using ordinary one-way ANOVA multiple comparisons and comparison made to vhs* values. ns, p > 0.05. *** p > 0.001. **** p < 0.0001.

To explore this relative activity further, we next carried out a dose response of each of these vhs-expressing plasmids in comparison to the vhs*-expressing plasmid and measured secreted GLuc activity. This indicated that as little as 10 ng of transfected vhsB-expressing plasmid reduced GLuc activity by over five-fold, greater than that seen for vhsH or vhsE (Fig 3B). A further dose response using even lower amounts of vhs plasmid (2 ng to 12 ng) indicated that as little as 2ng of vhsB plasmid was able to reduce GLuc activity by over 70% (Fig 3C). The vhsB homologue was reproducibly more powerful at reducing GLuc expression than vhsH or vhsE homologues under the conditions employed in this assay. Given that vhsB is expressed at higher levels than the other homologues (Fig 2A) this enhanced activity is likely to be due in part to higher levels of the vhsB protein in the cytoplasm of the cell.

The known effect of vhsH on cellular mRNA is as an endoribonuclease, and hence it is predicted that it’s activity on GLuc expression would be a consequence of enhanced decay of the GLuc mRNA. To determine the corresponding effect of the vhs homologues on the level of GLuc mRNA, transfections were carried out as above using a single dose of plasmid expressing untagged vhs, and the relative amount of GLuc mRNA was measured using RT-qPCR. This revealed that those vhs homologues that had greatly reduced GLuc expression at the protein level also had reduced the level of the GLuc transcript (Fig 3D).

### Differential effect of vhs homologues on PABPC1 relocalisation to the nucleus

The expression of vhs is known to cause a change to the steady state localisation of the PABPC1 protein from cytoplasmic to nuclear, an activity that is proposed to reflect the enhanced turnover of mRNA in the cytoplasm and the concomitant release of PABPC1 to recycle to the nucleus (23, 46). To test the ability of the vhs homologues to alter the compartmentalisation of PABPC1, we examined PABPC1 localisation in cells expressing untagged vhs homologues, which indicated that PABPC1 was found in the nucleus of up to 60% of the cell monolayer, with the percentage of cells containing nuclear PABPC1 broadly correlating with relative vhs activity: vhsB > vhsE > vhsH > vhsV (Fig 4A & 4B). The one exception to this gradient was vhsM, which induced PABPC1 relocalisation to the nucleus of over 30% of cells despite causing only a modest reduction in GLuc activity (Fig 4A &4B, vhsM). Strikingly, the predominant effect of vhsV expression was not to relocalise PABPC1 to the nucleus but rather to induce PABPC1-containing cytoplasmic accumulations (Fig 4A, 4B & 4C, vhsV), an outcome that was also present in combination with nuclear PABPC1 in vhsM expressing-cells (Fig 4A & 4C, vhsM).

**Figure 4.**
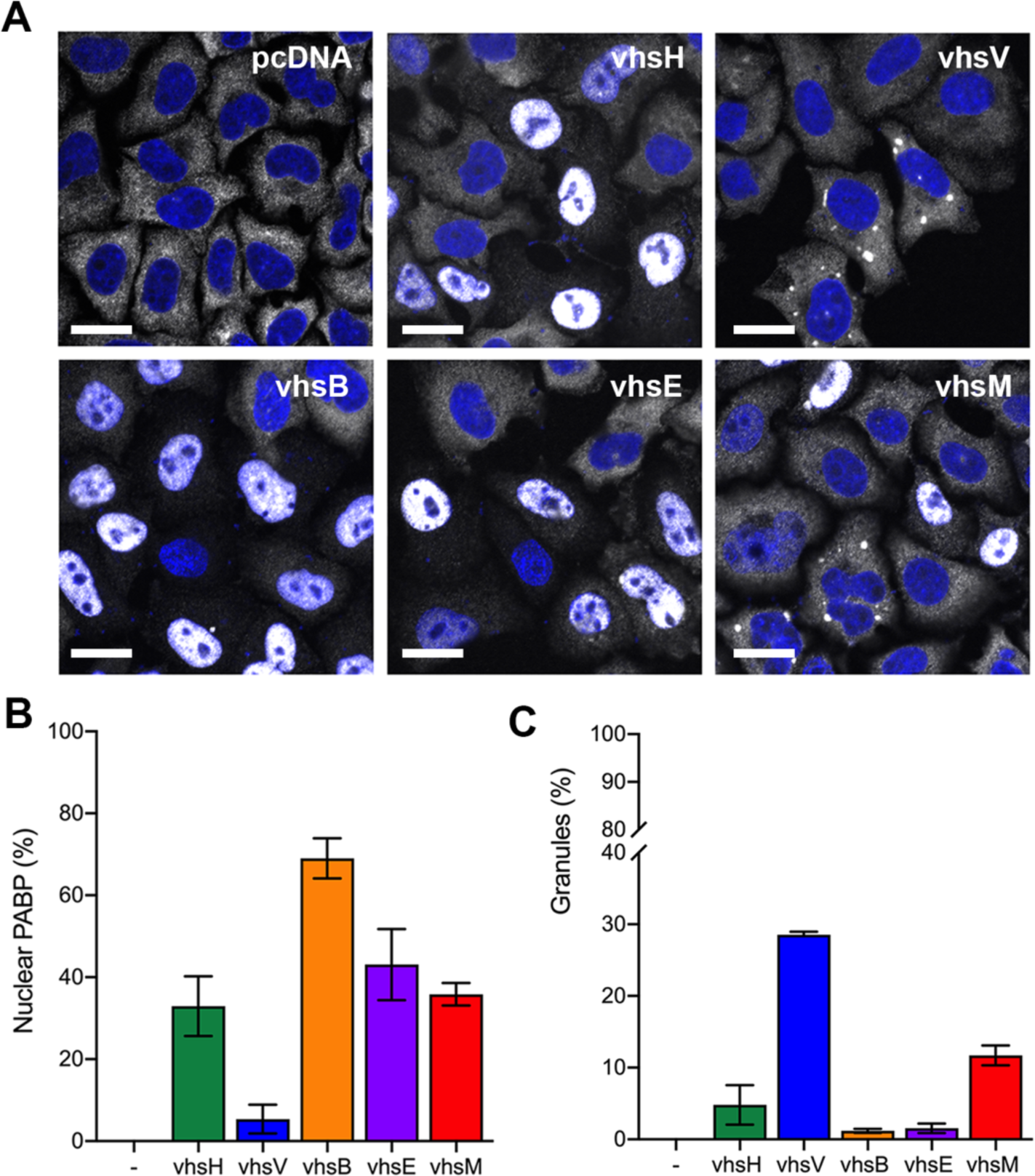
Differential effect of vhs homologue expression on PABPC1 localisation. **(A)** HeLa cells grown on coverslips were transfected with plasmids expressing untagged vhs homologues, fixed twenty hours later and stained for PABPC1 (white). Nuclei were stained with DAPI (blue) and cells were imaged on a NikonA1 confocal microscope. Scale bar = 20 μm. **(B)** Cells with nuclear PABPC1 shown in (B) were counted in three independent experiments (> 150 cells in each). (**C)** Cells with cytoplasmic granules of PABPC1 in (B) were counted in two independent experiments (> 150 cells in each).

### Vhs homologues that don’t induce mRNA decay induce the assembly of cytoplasmic G3BP1-containing RNA granules

To determine the effect of vhs homologue expression on global mRNA, we utilised *in situ* hybridisation of polyadenylated RNA using a fluorescently labelled oligodT probe. Cells incubated with oligodT were further stained for PABPC1 to correlate the global localisation of polyA tails with the PABPC1 polyA tail-binding protein (Fig 5). This revealed that PABPC1 localisation correlated with that of polyA+ RNA, with vhsH, vhsB and vhsE inducing the cytoplasmic depletion and the nuclear accumulation of the polyA+ signal. However, in the case of vhsV and vhsM, polyA+ RNA accumulated in cytoplasmic granules. These RNA granules resemble stress granules (SGs), which are membraneless biocondensates formed when translation is stalled, and contain preinitiation translation complexes, 40s ribosomal units and over 400 proteins including PAPBC1 and the classical SG marker, G3BP1 (47). We therefore used G3BP1 alongside PABPC1 in cells expressing each of the vhs homologues to show that the RNA granules induced by expression of vhsV and vhsM also contained G3BP1, suggesting that they are formed as a consequence of stalled translation (Fig 6). Furthermore, expression of vhsM and vhsV but not the other homologues also resulted in enhanced levels of both PKR and eIF2α phosphorylation suggesting that the RNA granules in these cells were a consequence of stress responses (Fig 7A & 7B).

**Figure 5.**
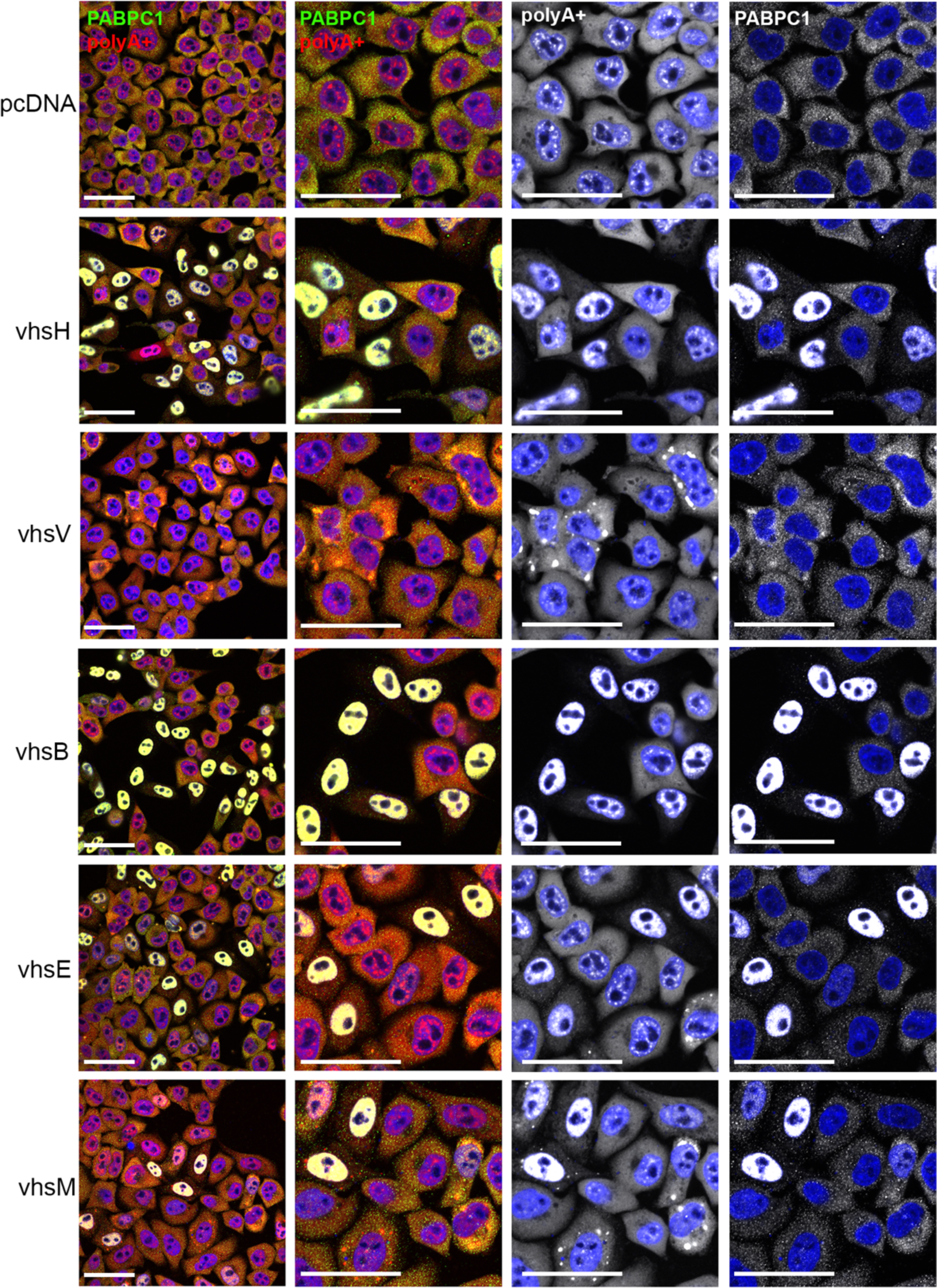
Differential effect of vhs homologues on the localisation of polyadenylated RNA. HeLa cells grown on coverslips were transfected with plasmids expressing untagged vhs homologues, fixed twenty hours later and processed first for polyA+ FISH (red) and second for PABPC1 immunofluorescence (green). Nuclei were stained with DAPI (blue) and cells were imaged on a NikonA1 confocal microscope. Scale bar = 50 μm.

**Figure 6:**
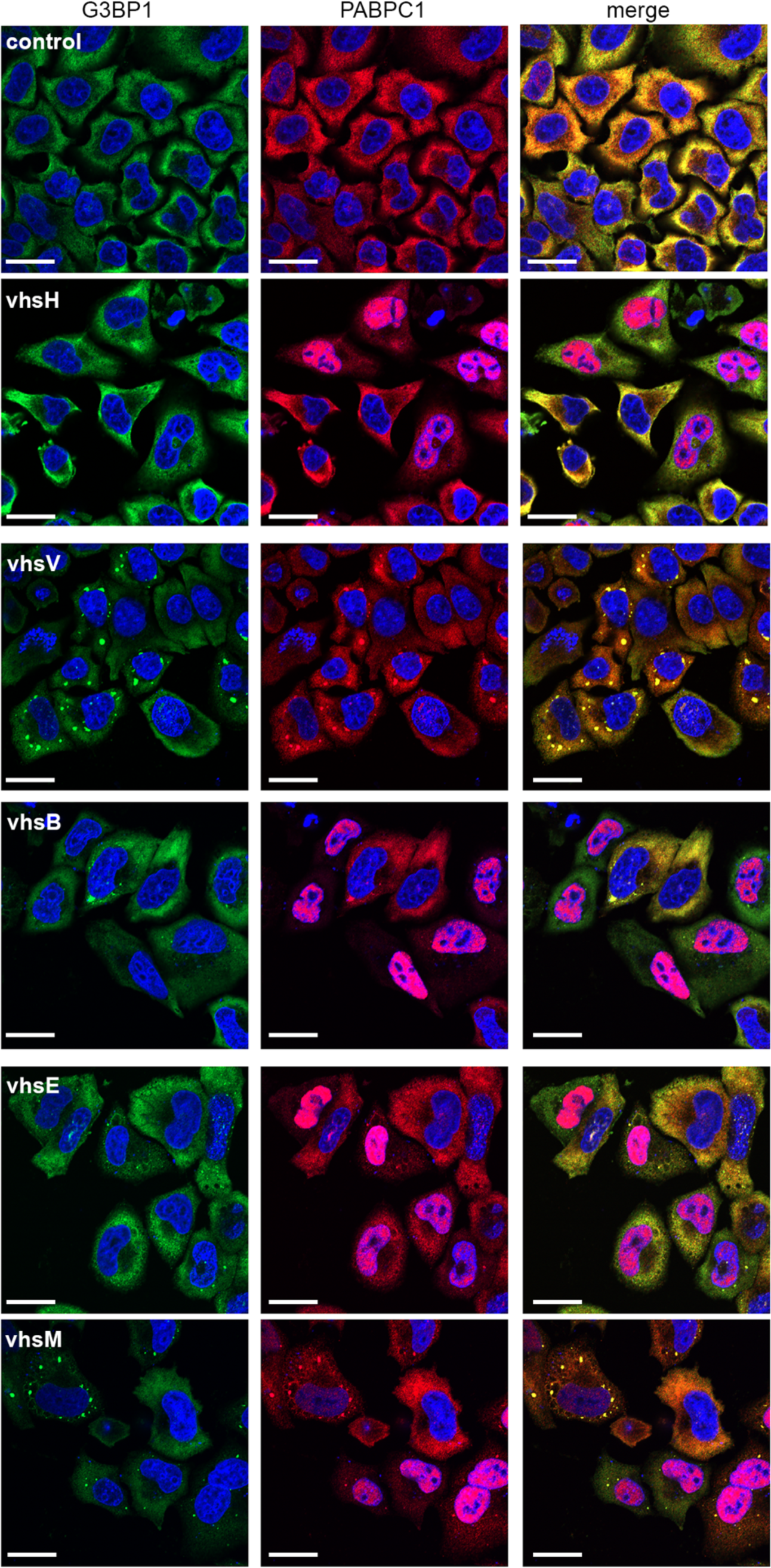
VZV and MDV homologues of vhs induce assembly of G3BP1-containing RNA granules. HeLa cells grown on coverslips were transfected with plasmids expressing vhs homologues, fixed twenty hours later and stained for G3BP1 (green) and PABPC1 (red). Nuclei were stained with DAPI (blue) and cells were imaged on a NikonA1 confocal microscope. Scale bar = 20 μm.

**Figure 7.**
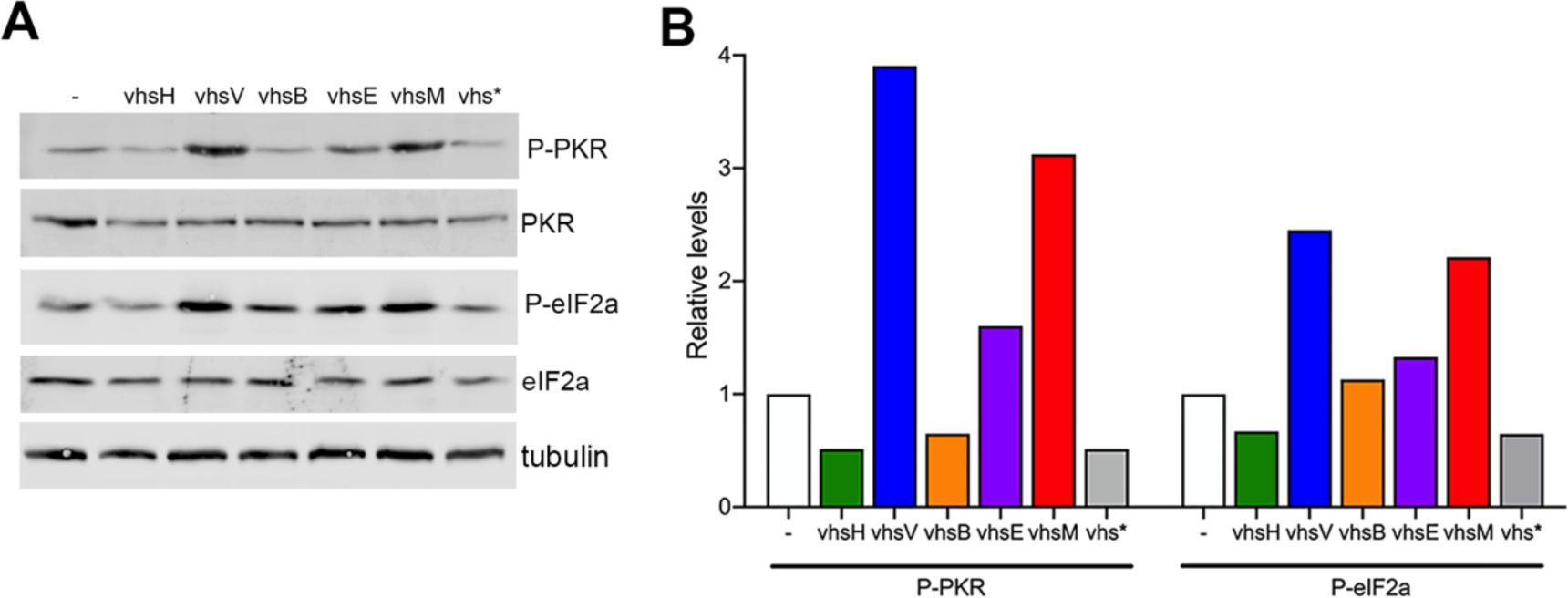
Induction of PKR and eIF2α phosphorylation correlates with stress granule formation by vhsV and vhsM. **(A)** Lysates of HeLa cells transfected with plasmids expressing each of the vhs homologues were analysed by SDS-PAGE and Western blotting with the indicated antibodies. **(B**) Quantification of the Western blots shown in A using LI-COR Odyssey software. Phospho -PKR and -eIF2α were normalised to the total of each protein and subsequently to α-tubulin.

### All homologues of vhs induce global translational arrest

To measure translational arrest directly, we next carried out single-cell puromycin incorporation assays of HeLa cells expressing each of the untagged vhs homologues, and subsequently stained for both puromycin together with PABPC1, used here as a surrogate for vhs expression (Fig. 8). This indicated that in cells expressing vhsH, vhsB or vhsE, puromycin incorporation was greatly reduced in cells where PABPC1 was nuclear and hence where mRNA decay had occurred. However, in cells expressing vhsV or vhsM where cytoplasmic PABPC1 granules had formed, puromycin incorporation was also reduced to a similar degree. Hence, the two vhs homologues that are shown above to be defective at inducing mRNA degradation, instead appear to induce translational arrest through the formation of RNA granules.

**Figure 8:**
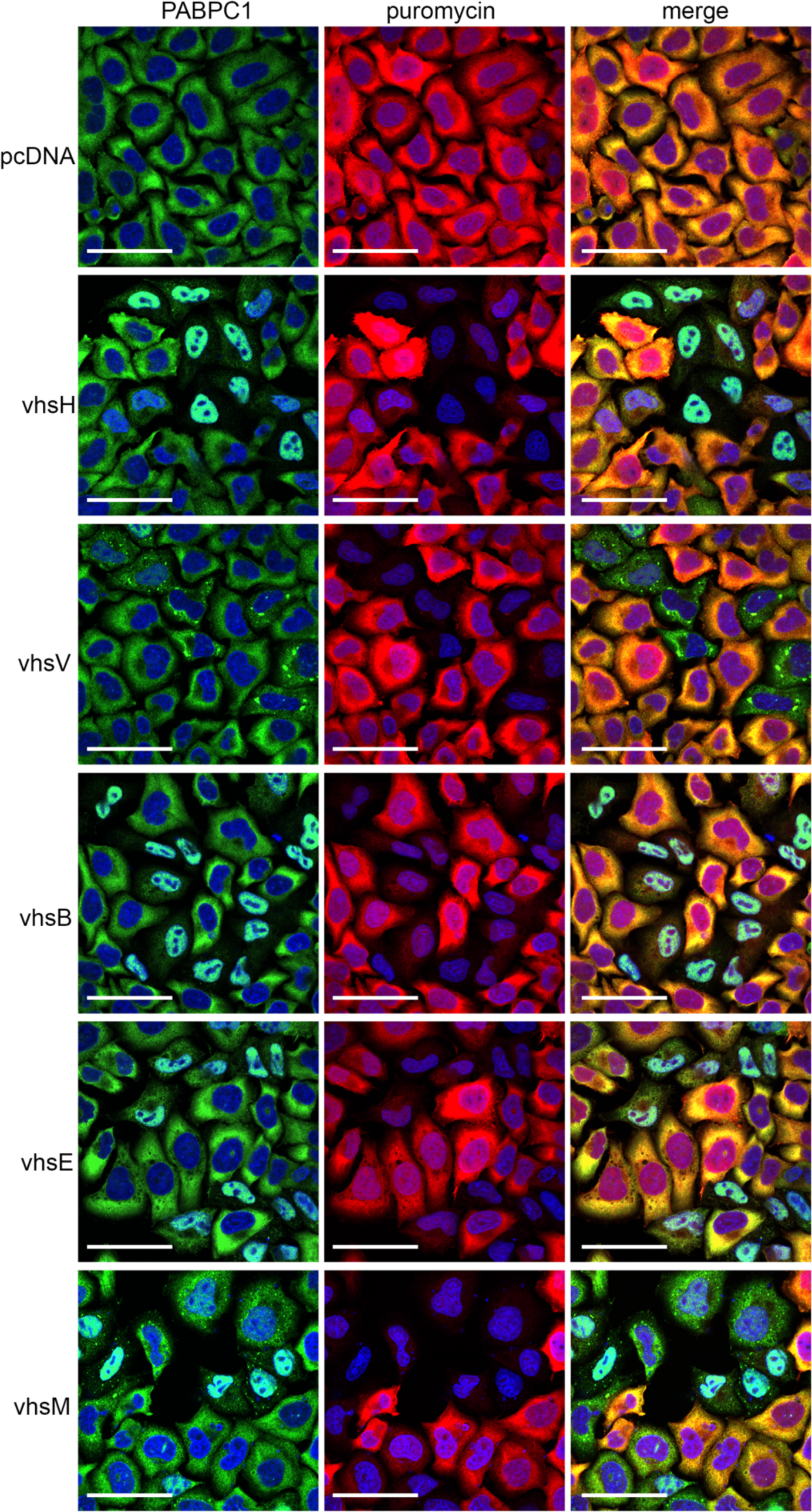
All vhs homologues induce translational blockade regardless of mRNA decay activity. HeLa cells grown on coverslips were transfected with plasmids expressing each of the vhs homologues. After 16h, cells were incubated with puromycin for 10 mins before fixation and permeabilization. Cells were then stained with antibodies to PABPC1 (green) and puromycin (red), nuclei stained with DAPI (blue) and images acquired using a Nikon A1 confocal microscope. Scale bar = 50 μm.

## Discussion

In this study we have utilised a comparative biology approach to refine the functional pathway of the conserved alphaherpesvirus protein vhs, which in HSV1 has been shown to be an endoribonuclease. Using a combination of reporter assays and single cell biological studies, we have shown that the vhs homologues fall into two groups: those that induce the degradation of mRNA resulting in translational shutoff, and those that only cause a block to translation. These unexpected results not only contribute to the ongoing molecular elucidation of vhs activity, but also help to unravel the potential role and mechanism of vhs in alphaherpesvirus infections other than HSV1.

It has generally been accepted that translational shutoff in HSV1 infection is a consequence of the global mRNA decay induced by vhsH. Indeed, the data presented here and elsewhere on ectopic expression of vhsH support that global RNA decay and translational shutoff go hand-in-hand in cells expressing not only vhsH, but also vhsB and vhsE. Nonetheless, vhsH does not cleave RNA indiscriminately but rather has been shown to be specific for mRNA, a mechanism that has been proposed to occur by binding to the translational initiation machinery at the 5’ end of the mRNA (18–21). As such, this provides scope for vhsH protein to shut down translation prior to and without the need for mRNA decay. Intriguingly, the ectopic expression of the vhsV homologue does not induce either decay of the GLuc reporter mRNA or the cytoplasmic depletion of polyadenylated RNA, yet blocks global protein synthesis as measured by greatly reduced puromycin incorporation, as well as a significant reduction in GLuc activity. This suggests that vhsV, which is encoded by the ORF17 gene of VZV, retains the capacity to block translation by interaction with the mRNA CAP-binding complex but does not possess endoribonuclease activity - at least when expressed in isolation of virus infection - a result that may in some way explain the discrepancies in previous studies of vhsV ORF17 functionality (42, 43). Consequently, vhsB and vhsV appear to sit at two ends of the vhs endoribonuclease activity spectrum, exhibiting the greatest and least effect on polyadenylated RNA/PABPC1 relocalisation respectively. Intriguingly, vhsM exhibited a mixed phenotype, such that when expressed in HeLa cells, it had similar characteristics to vhsV, as it was unable to degrade GLuc mRNA but at least in some cells PABPC1 was relocalised to the nucleus. However, MDV is a virus that infects chickens, so it is likely that vhsM cannot function correctly in mammalian cells due to differences in important cellular factors. Hence, further studies in chicken cells will be needed to further define the activity of vhsM in an appropriate system. Taken together, this leads us to propose that there are two steps to vhs activity: 1) binding to the translational initiation machinery leading to translational blockade, and 2) the cleavage of associated mRNA leading to global mRNA decay; all homologues possess the ability to undertake step 1, but vhsV and to a large extent vhsM are defective in step 2.

It has been shown before that infection with HSV1 or HSV2 lacking vhs induces the assembly of stress granules (48, 49). It has also been proposed that vhs is required to degrade dsRNA formed in infected cells, and in its absence, dsRNA is bound by PKR which is activated to phosphorylate eIF2α and inhibit translation initiation. The downstream outcome of this would be the assembly of stress granules in Δvhs infected cells. Given this proposed role for vhs in blocking stress granule assembly, it is therefore counterintuitive that the activity of vhsV and vhsM homologues in fact induce the assembly of RNA granules similar to canonical stress granules and enhance the phosphorylation of PKR and eIF2α, rather than degrade mRNA. Although PKR phosphorylation of the translation initiation factor eIF2α typically induces translational arrest and subsequent stress granule assembly, stress granules themselves have been shown to activate PKR and subsequent eIF2α phosphorylation (50, 51), raising the possibility that this is how vhsV and vhsM inhibit translation within the cell.

One of the defining characteristics of vhsH is that, when introduced ectopically on a plasmid, it is expressed at very low levels (13). Moreover, this low level of expression is maintained when vhsH is expressed as a GFP fusion protein (44), suggesting that it is an inherent feature of the vhsH ORF. We have previously described three reasons for this low level of vhs protein: first, because vhsH is an endoribonuclease, it can induce the degradation of its own transcript, thereby inducing a negative feedback loop to control its expression (13). Second, we have also shown that unlike other transcripts which accumulate in the cytoplasm, the vhsH transcript is inherently nuclear, providing an additional mechanism for regulation of vhs expression (13), and as shown here, this property is conserved across all the vhs homologues tested. Third, we have also identified a short region of the vhsH ORF known as the inhibitory sequence (IS) which is fully transferable to other transcripts and reduces their expression (13). It is therefore noteworthy that all homologues of vhs tested here also conferred a similarly low level of GFP expression when fused to the GFP ORF, suggesting an equivalent mechanism of autoregulation amongst the vhs homologues. Whether any or all of these homologues contain translation inhibitory sequences similar to that found in vhsH remains to be determined, but there was a gradient of expression of vhsB>vhsE>vhsM>vhsH>vhsV. The relatively high expression of vhsB in comparison to vhsH may partially explain its enhanced activity in the assays we have utilised but this is yet to be confirmed.

The data presented here has allowed us to refine our overarching model for vhs activity. In this model, all vhs homologues tested would bind to mRNAs via the translation initiation machinery, resulting in translational stalling. In the absence of mRNA decay, translational stalling would lead to the assembly of cytoplasmic RNA granules, an amplification of the PKR response pathway, and a global shutdown of protein synthesis. However, endonuclease active vhs homologues would cleave associated mRNAs thereby inducing their degradation and release of PABPC1 to the nucleus. This mRNA degradation would block the assembly of RNA granules induced by vhs, freeing up ribosomal subunits and relevant RBPs for further activity. While there are many steps of this pathway that still need to be defined, these data reveal the power of a comparative biology approach to understand not only how vhs functions but also key steps in the regulation of cellular mRNA decay and translational control.

## Acknowledgments

The authors thank Dr Yongxiu Yao (The Pirbright Institute) for MDV strain RB1B genomic DNA, and Dr Neil Bryant (Animal Health Trust) for EHV1 strain Ab4 genomic DNA.

## References

1. Gaucherand L, Porter BK, Levene RE, Price EL, Schmaling SK, Rycroft CH, Kevorkian Y, McCormick C, Khaperskyy DA, Gaglia MM. 2019. The Influenza A Virus Endoribonuclease PA-X Usurps Host mRNA Processing Machinery to Limit Host Gene Expression. Cell Rep 27:776–792 e7.

2. Jagger BW, Wise HM, Kash JC, Walters KA, Wills NM, Xiao YL, Dunfee RL, Schwartzman LM, Ozinsky A, Bell GL, Dalton RM, Lo A, Efstathiou S, Atkins JF, Firth AE, Taubenberger JK, Digard P. 2012. An overlapping protein-coding region in influenza A virus segment 3 modulates the host response. Science 337:199–204.

3. Glaunsinger B, Ganem D. 2004. Lytic KSHV infection inhibits host gene expression by accelerating global mRNA turnover. Mol Cell 13:713–23.

4. Read GS. 2013. Virus-encoded endonucleases: expected and novel functions. Wiley Interdiscip Rev RNA 4:693–708.

5. Bhardwaj K, Guarino L, Kao CC. 2004. The severe acute respiratory syndrome coronavirus Nsp15 protein is an endoribonuclease that prefers manganese as a cofactor. J Virol 78:12218–24.

6. Covarrubias S, Richner JM, Clyde K, Lee YJ, Glaunsinger BA. 2009. Host shutoff is a conserved phenotype of gammaherpesvirus infection and is orchestrated exclusively from the cytoplasm. J Virol 83:9554–66.

7. Read GS, Frenkel N. 1983. Herpes simplex virus mutants defective in the virion-associated shutoff of host polypeptide synthesis and exhibiting abnormal synthesis of alpha (immediate early) viral polypeptides. J Virol 46:498–512.

8. Kwong AD, Frenkel N. 1989. The herpes simplex virus virion host shutoff function. J Virol 63:4834–9.

9. Read GS, Karr BM, Knight K. 1993. Isolation of a herpes simplex virus type 1 mutant with a deletion in the virion host shutoff gene and identification of multiple forms of the vhs (UL41) polypeptide. J Virol 67:7149–60.

10. Matis J, Kudelova M. 2001. Early shutoff of host protein synthesis in cells infected with herpes simplex viruses. Acta Virol 45:269–77.

11. Jones FE, Smibert CA, Smiley JR. 1995. Mutational analysis of the herpes simplex virus virion host shutoff protein: evidence that vhs functions in the absence of other viral proteins. J Virol 69:4863–71.

12. Everly DN, Jr., Read GS. 1997. Mutational analysis of the virion host shutoff gene (UL41) of herpes simplex virus (HSV): characterization of HSV type 1 (HSV-1)/HSV-2 chimeras. J Virol 71:7157–66.

13. Elliott G, Pheasant K, Ebert-Keel K, Stylianou J, Franklyn A, Jones J. 2018. Multiple post-transcriptional strategies to regulate the herpes simplex virus type 1 vhs endoribonuclease. J Virol doi:10.1128/JVI.00818-18.

14. Strand SS, Leib DA. 2004. Role of the VP16-binding domain of vhs in viral growth, host shutoff activity, and pathogenesis. J Virol 78:13562–72.

15. Oroskar AA, Read GS. 1989. Control of mRNA stability by the virion host shutoff function of herpes simplex virus. J Virol 63:1897–906.

16. Elgadi MM, Hayes CE, Smiley JR. 1999. The herpes simplex virus vhs protein induces endoribonucleolytic cleavage of target RNAs in cell extracts. J Virol 73:7153–64.

17. Taddeo B, Roizman B. 2006. The virion host shutoff protein (UL41) of herpes simplex virus 1 is an endoribonuclease with a substrate specificity similar to that of RNase A. J Virol 80:9341–5.

18. Everly DN, Jr., Feng P, Mian IS, Read GS. 2002. mRNA degradation by the virion host shutoff (Vhs) protein of herpes simplex virus: genetic and biochemical evidence that Vhs is a nuclease. J Virol 76:8560–71.

19. Feng P, Everly DN, Jr., Read GS. 2005. mRNA decay during herpes simplex virus (HSV) infections: protein-protein interactions involving the HSV virion host shutoff protein and translation factors eIF4H and eIF4A. J Virol 79:9651–64.

20. Page HG, Read GS. 2010. The virion host shutoff endonuclease (UL41) of herpes simplex virus interacts with the cellular cap-binding complex eIF4F. J Virol 84:6886–90.

21. Doepker RC, Hsu WL, Saffran HA, Smiley JR. 2004. Herpes simplex virus virion host shutoff protein is stimulated by translation initiation factors eIF4B and eIF4H. J Virol 78:4684–99.

22. Covarrubias S, Gaglia MM, Kumar GR, Wong W, Jackson AO, Glaunsinger BA. 2011. Coordinated destruction of cellular messages in translation complexes by the gammaherpesvirus host shutoff factor and the mammalian exonuclease Xrn1. PLoS Pathog 7:e1002339.

23. Kumar GR, Glaunsinger BA. 2010. Nuclear import of cytoplasmic poly(A) binding protein restricts gene expression via hyperadenylation and nuclear retention of mRNA. Mol Cell Biol 30:4996–5008.

24. Pheasant K, Moller-Levet CS, Jones J, Depledge D, Breuer J, Elliott G. 2018. Nuclear-cytoplasmic compartmentalization of the herpes simplex virus 1 infected cell transcriptome is co-ordinated by the viral endoribonuclease vhs and cofactors to facilitate the translation of late proteins. PLoS Pathog 14:e1007331.

25. Afonina E, Stauber R, Pavlakis GN. 1998. The human poly(A)-binding protein 1 shuttles between the nucleus and the cytoplasm. J Biol Chem 273:13015–21.

26. Burgess HM, Richardson WA, Anderson RC, Salaun C, Graham SV, Gray NK. 2011. Nuclear relocalisation of cytoplasmic poly(A)-binding proteins PABP1 and PABP4 in response to UV irradiation reveals mRNA-dependent export of metazoan PABPs. J Cell Sci 124:3344–55.

27. Gilbertson S, Federspiel JD, Hartenian E, Cristea IM, Glaunsinger B. 2018. Changes in mRNA abundance drive shuttling of RNA binding proteins, linking cytoplasmic RNA degradation to transcription. Elife 7.

28. Dauber B, Saffran HA, Smiley JR. 2019. The herpes simplex virus host shutoff (vhs) RNase limits accumulation of double stranded RNA in infected cells: Evidence for accelerated decay of duplex RNA. PLoS Pathog 15:e1008111.

29. Dauber B, Pelletier J, Smiley JR. 2011. The herpes simplex virus 1 vhs protein enhances translation of viral true late mRNAs and virus production in a cell type-dependent manner. J Virol 85:5363–73.

30. Esclatine A, Taddeo B, Roizman B. 2004. Herpes simplex virus 1 induces cytoplasmic accumulation of TIA-1/TIAR and both synthesis and cytoplasmic accumulation of tristetraprolin, two cellular proteins that bind and destabilize AU-rich RNAs. J Virol 78:8582–92.

31. Anderson P, Kedersha N. 2006. RNA granules. J Cell Biol 172:803–8.

32. Berthomme H, Jacquemont B, Epstein A. 1993. The pseudorabies virus host-shutoff homolog gene: nucleotide sequence and comparison with alphaherpesvirus protein counterparts. Virology 193:1028–32.

33. Everly DN, Jr., Read GS. 1999. Site-directed mutagenesis of the virion host shutoff gene (UL41) of herpes simplex virus (HSV): analysis of functional differences between HSV type 1 (HSV-1) and HSV-2 alleles. J Virol 73:9117–29.

34. Lin HW, Chang YY, Wong ML, Lin JW, Chang TJ. 2004. Functional analysis of virion host shutoff protein of pseudorabies virus. Virology 324:412–8.

35. Ma W, Wang H, He H. 2019. Bovine herpesvirus 1 tegument protein UL41 suppresses antiviral innate immune response via directly targeting STAT1. Vet Microbiol 239:108494.

36. Hinkley S, Ambagala AP, Jones CJ, Srikumaran S. 2000. A vhs-like activity of bovine herpesvirus-1. Arch Virol 145:2027–46.

37. Dai H, Wu J, Yang H, Guo Y, Di H, Gao M, Wang J. 2022. Construction of BHV-1 UL41 Defective Virus Using the CRISPR/Cas9 System and Analysis of Viral Replication Properties. Front Cell Infect Microbiol 12:942987.

38. Lin HW, Hsu WL, Chang YY, Jan MS, Wong ML, Chang TJ. 2010. Role of the UL41 protein of pseudorabies virus in host shutoff, pathogenesis and induction of TNF-alpha expression. J Vet Med Sci 72:1179–87.

39. Gimeno I, Silva RF. 2008. Deletion of the Marek’s disease virus UL41 gene (vhs) has no measurable effect on latency or pathogenesis. Virus Genes 36:499–507.

40. Mao W, Niikura M, Silva RF, Cheng HH. 2008. Quantitative evaluation of viral fitness due to a single nucleotide polymorphism in the Marek’s disease virus UL41 gene via an in vitro competition assay. J Virol Methods 148:125–31.

41. Feng X, Thompson YG, Lewis JB, Caughman GB. 1996. Expression and function of the equine herpesvirus 1 virion-associated host shutoff homolog. J Virol 70:8710–8.

42. Sato H, Callanan LD, Pesnicak L, Krogmann T, Cohen JI. 2002. Varicella-zoster virus (VZV) ORF17 protein induces RNA cleavage and is critical for replication of VZV at 37 degrees C but not 33 degrees C. J Virol 76:11012–23.

43. Desloges N, Rahaus M, Wolff MH. 2005. The varicella-zoster virus-mediated delayed host shutoff: open reading frame 17 has no major function, whereas immediate-early 63 protein represses heterologous gene expression. Microbes Infect 7:1519–29.

44. Wise EL, Samolej J, Elliott G. 2022. Herpes Simplex Virus 1 Expressing GFP-Tagged Virion Host Shutoff (vhs) Protein Uncouples the Activities of RNA Degradation and Differential Nuclear Retention of the Virus Transcriptome. J Virol doi:10.1128/jvi.01926-21:e0192621.

45. Taddeo B, Sciortino MT, Zhang W, Roizman B. 2007. Interaction of herpes simplex virus RNase with VP16 and VP22 is required for the accumulation of the protein but not for accumulation of mRNA. Proc Natl Acad Sci U S A 104:12163–8.

46. Burgess HM, Gray NK. 2012. An integrated model for the nucleo-cytoplasmic transport of cytoplasmic poly(A)-binding proteins. Commun Integr Biol 5:243–7.

47. Kedersha N, Stoecklin G, Ayodele M, Yacono P, Lykke-Andersen J, Fritzler MJ, Scheuner D, Kaufman RJ, Golan DE, Anderson P. 2005. Stress granules and processing bodies are dynamically linked sites of mRNP remodeling. J Cell Biol 169:871–84.

48. Finnen RL, Hay TJ, Dauber B, Smiley JR, Banfield BW. 2014. The herpes simplex virus 2 virion-associated ribonuclease vhs interferes with stress granule formation. J Virol 88:12727–39.

49. Dauber B, Poon D, Dos Santos T, Duguay BA, Mehta N, Saffran HA, Smiley JR. 2016. The Herpes Simplex Virus Virion Host Shutoff Protein Enhances Translation of Viral True Late mRNAs Independently of Suppressing Protein Kinase R and Stress Granule Formation. J Virol 90:6049–6057.

50. Burgess HM, Mohr I. 2018. Defining the role of stress granules in innate immune suppression by the HSV-1 endoribonuclease VHS. J Virol doi:10.1128/JVI.00829-18.

51. Reineke LC, Kedersha N, Langereis MA, van Kuppeveld FJ, Lloyd RE. 2015. Stress granules regulate double-stranded RNA-dependent protein kinase activation through a complex containing G3BP1 and Caprin1. mBio 6:e02486.

